# Multi-Omics and Machine Learning-Based Profiling of Severity Signatures in *Mycoplasma Pneumoniae* Infection in Children

**DOI:** 10.1101/2025.04.05.647347

**Authors:** Guiqiu Li, Wenzheng Wang, Zhili Hu, Qiaowen Yang, Lili He, Yun Gao, Xiulan Lai

## Abstract

Mycoplasma pneumoniae pneumonia (MPP) is a common respiratory infection in children, yet the mechanisms driving its progression to severe disease remain poorly understood. This study employs a comprehensive proteomic and metabolomic approach to elucidate severity-related pathways and identify potential biomarkers for improved diagnosis and targeted therapy. By analyzing blood proteomes from 57 pediatric patients with varying MPP severities alongside 10 healthy controls, and integrating multi-omics data from bronchoalveolar lavage fluid (BALF), we uncovered key severity-associated proteins and metabolites linked to inflammatory and metabolic dysregulation. Notably, alterations in L-arginine metabolism, influenced by APAF1 and SERPINB5, were found to modulate the efferocytosis pathway, while ERCC1 and MLH1, crucial components of the Fanconi anemia pathway, were associated with changes in plasmid acid metabolites affecting fatty acid elongation. Machine learning analysis further identified three critical biomarkers—TNFRSF10B, SAT1, and 4-nitrophenol— that accurately distinguished between mild and severe MPP cases with high sensitivity (0.8) and specificity (1). These findings provide novel insights into the molecular mechanisms underlying severe MPP, highlighting efferocytosis and fatty acid metabolism as key pathways. The identification of severity-specific biomarkers offers a foundation for enhanced diagnostic precision, improved disease stratification, and the development of targeted therapeutic strategies to optimize the management of severe MPP in pediatric patients.

**Highlights:**

- Identified severity signatures in *Mycoplasma pneumoniae* infection in children.
- Conducted the multi-omics analysis of bronchoalveolar lavage fluid (BALF) in children with varying severities of *Mycoplasma pneumoniae* infection.
- Discovered biomarkers that effectively distinguish mild from severe *Mycoplasma pneumoniae* infection.
- Findings offer potential targets to improve the diagnosis and treatment of severe *Mycoplasma pneumoniae*
- infection in children.

## Introduction

*Mycoplasma Pneumoniae* (MPP) is a pathogen that lacks a cell wall and belongs to the Mycoplasmataceae family. It is a common cause of respiratory infections, particularly community-acquired pneumonia in children.^1,2^ The infection is primarily transmitted through respiratory droplets, especially in crowded envrieonments.^3^ MPP infections usually start with mild respiratory symptoms such as fever, cough, and sore throat. In some cases, the infection can spread and cause pneumonia.^4^ children is more vulnerable to infections caused by the agent due to the vulnerable immune system.^5–7^ It may start with mild upper respiratory symptoms caused by MPP infections. However, in severe cases, the infection can progress to bronchitis or pneumonia,which coud lead to difficulty breathing and a persistent high fever.^8,9^

Currently, Omics techniques are becoming indispensable in studying infectious diseases, providing insight into the molecular mechanisms underlying pathogen-host interactions, immune responses, and disease progression.^10,11^ Progress in proteomic and metabolomic analyses, and their application to challenge infectious diseases by discovering proteins in biofluids during infection, thus linking greater prominence and development of being valuable for early diagnosis, monitoring disease progression, and assessing therapeutic effectiveness.^12–14^ The application of proteomic and metabolics technologies allows for the comprehensive analysis of protein abundance, modifications, and interactions, enabling a deeper understanding of host-pathogen dynamics and facilitating the development of novel diagnostic and therapeutic strategies for infectious diseases.^15,16^ However, proteomic and metabolomic analyses of bronchoalveolar lavage fluid in patients infected with MPP remain limited. Therefore, analyzing protein-metabolite associations in this fluid can help pinpoint the specific causes of severe MPP infections in children.

In this study, we conducted a proteomic analysis of plasma samples from healthy children and children infected with MPP. The results indicated that the severity of MPP infection correlates with an inflammatory response. Further analysis was conducted on bronchoalveolar lavage fluid from MPP patients with different degrees of severity. This analysis, combined with metabolomic profiling, revealed a significant impact of MPP on inflammation. The affected pathways included p53 signaling and Fanconi anemia, involving proteins such as APAF1, SERPINB5, ERCC1, and MLH1. These inflammation-related pathways are likely to contribute to efferocytosis and fatty acid elongation, involving L-arginine and plasminogen acid, which are implicated in the pathogenesis of MPP. Furthermore, We used a random forest model to train on the differential proteins and metabolites and identified key biomarkers (TNFRSF10B, SAT1, and 4-nitrophenol). The models successfully distinguished between severe and mild cases of MPP infection (AUC = 0.9). In conclusion, this study highlights immune-related biomarkers involved in the pathogenesis of MPP and advances our understanding of MPP infections

## Results

### Clinical characteristics between the SMPP and MMPP

There is no intra-group bias in terms of age and gender between the SMPP and MMPP (p>0.05). In terms of clinical symptoms, body temperature, respiratory rate, and pleural effusion were comparable in both groups. However, it is of significance that all children were having cough. The laboratory examination revealed a significant difference in PCT and SAA between the two groups (P < 0.05), while no substantial difference was found in blood routine, CRP, IL-6, or D-dimer. The length of hospitalization for the children in both groups ranged from 5 to 9 days without significant variation.

### Blood proteome profiles for MPP infection patients and healthy controls

PCA analysis clearly separated plasma samples from mild, severe, and healthy individuals, suggesting that their plasma proteome profiles were significantly different (Figure 2A). Volcano plots produced from other targeted metabolomics analyses regulated those metabolites that were up (from the healthy, severe, and mild individuals - red) or down (blue) in the plasma (Figure 2B). Following the Mfuzz cluster analysis of protein expression, the clusters 5 and 7 patterns showed that the expression trends might correlate with the severity of the disease (Figure 2C). Overlapping differentially expressed proteins, with those occurring in clusters 5 and 7, were selected for a pathogenic analysis (Figure 2D). In an effort to better understand the underlying biological processes, we performed KEGG pathway enrichment and GO functional analyses on the intersection of proteins within cluster and the differentially expressed proteins (Figures 2E, S1). Interestingly, in MPPs the most expressed proteins were concentrated in the complement and coagulation cascades signaling pathway (Figure 2E).

**Figure 1.**
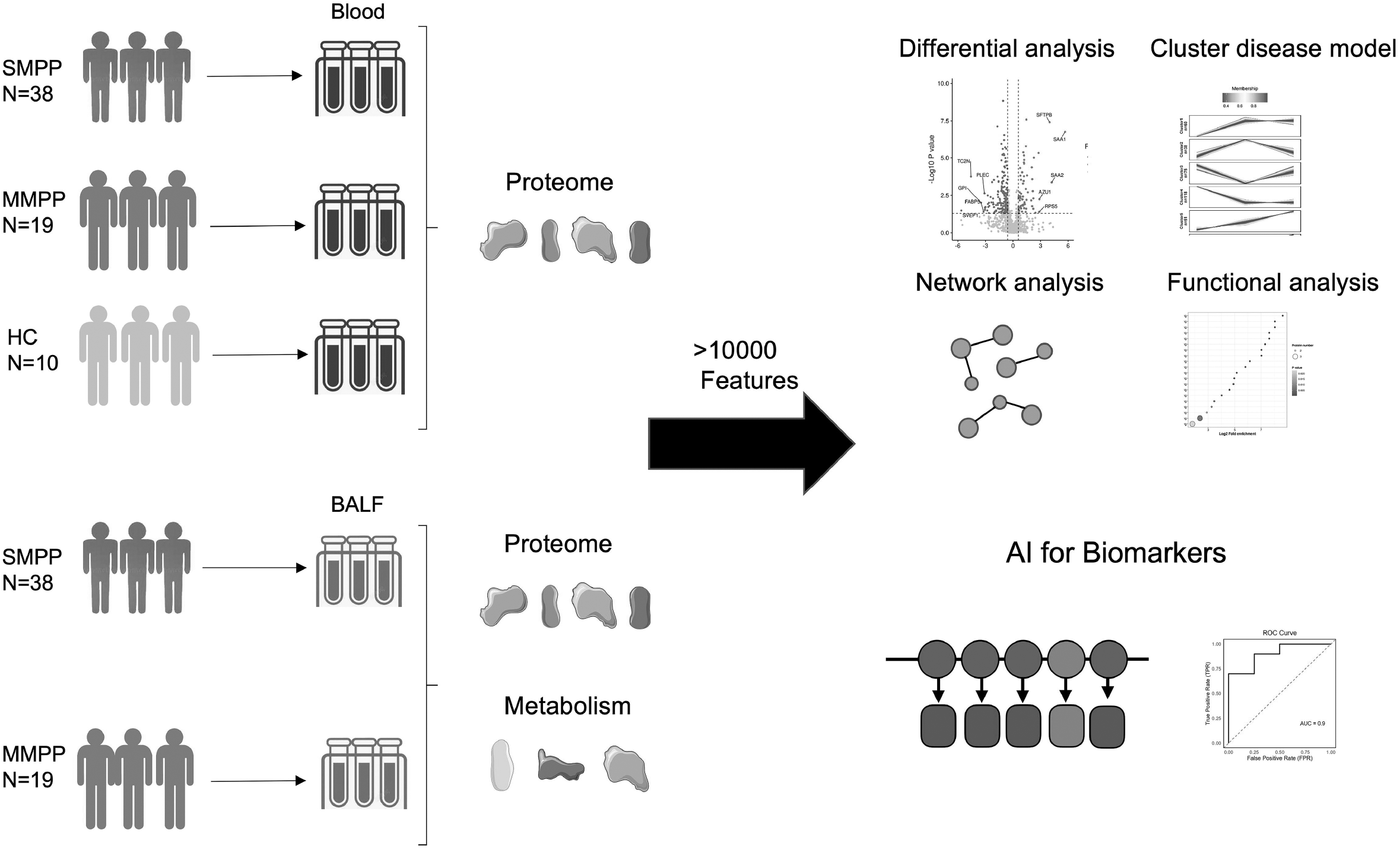
Investigation Framework for Omics-Based Analysis of MPP Patients and Healthy Individuals.

**Figure 2.**
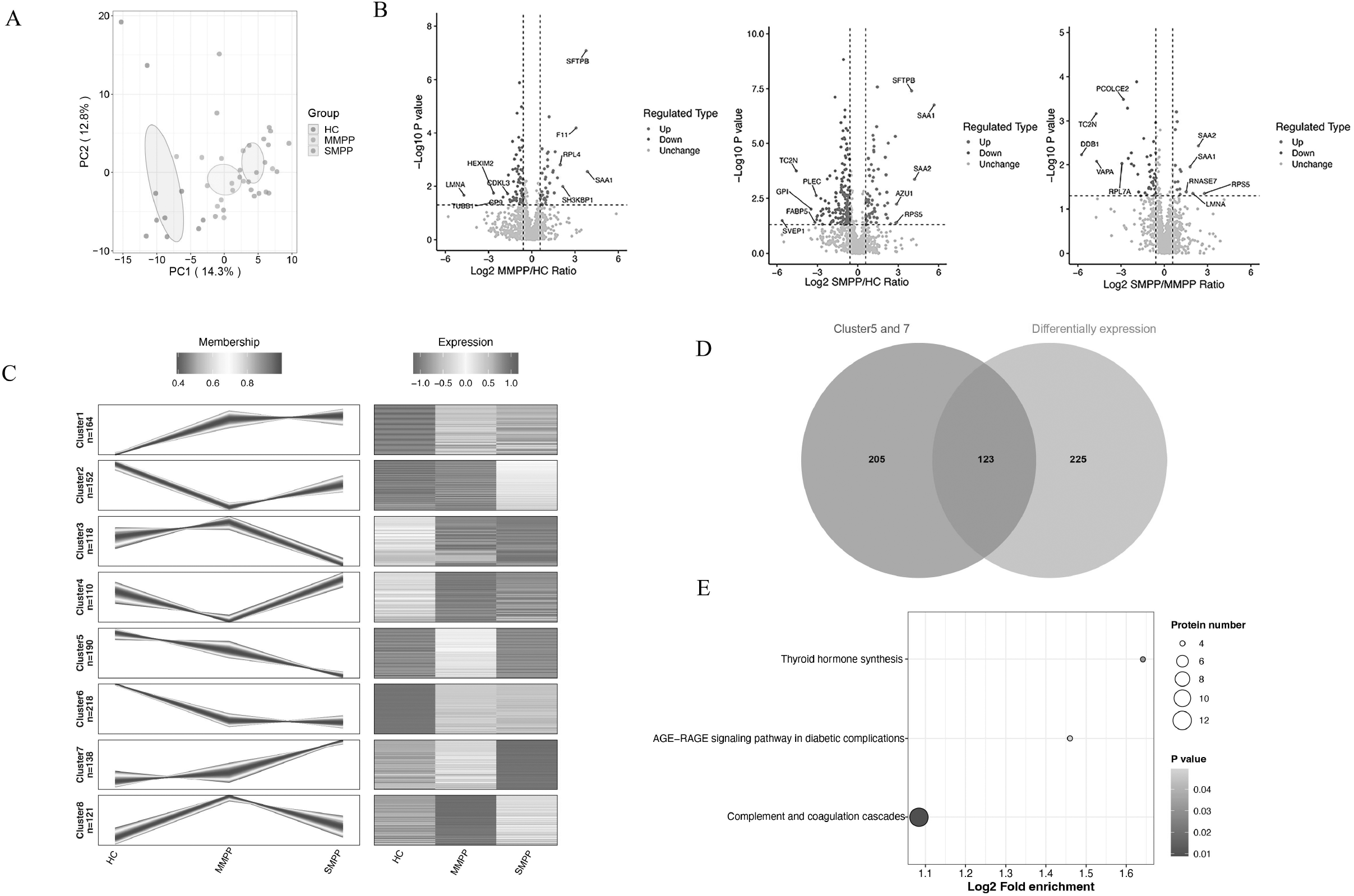
Proteomics Analysis between severe and non-severe patients in blood. A. The proteomics signatures analysis by Principal Component Analysis (PCA). B. The volcano plot derived from a targeted metabolomic analysis illustrates the top BALF proteomics that were increased (shown in red) or decreased (shown in blue) for the HC, MMPP, and SMPP groups. C. Clustering of proteins expression patterns across HC, MMPP, and SMPP groups. D. Venn diagram analysis of differential proteins and expression pattern clustering analysis. E. KEGG pathway enrichment analysis of overlapping proteins.

### BALF proteome profiles for patients with MPP infection

To examine for more pronounced differences between severe and mild groups, bronchoalveolar lavage fluid (BALF) samples were collected from 57 patients with differing severities for analysis. PCA analysis showed that the samples from severe patients and mild patients were nicely clustered apart, which illustrated significant differences in the profile of BALF proteomes (Figure 3A). Volcano plots from the untargeted proteomics analyses actually marked the metabolites that were found to increase (in red) or decrease (in blue) associated with severe and mild patients, respectively (Figure 3B). GO functional and KEGG pathway enrichment analysis was carried out on the intersection of the proteome (Figure 3C, S2). The most differentially expressed proteomes in MPPs were also highly concentrated in pathways related to inflammation, such as arginine and proline metabolism, the Fanconi anemia pathway, and the p53 signaling pathway. Furthermore, we conducted a PPI network analysis on 81 enriched proteins. Among these were 20 proteins representing four major pathways, such as the Fanconi pathway, centered around ERCC1, with proteins like FANCI, ATRIP, TOP3A, MLH1, CKMT1A, and the p53 signaling pathway centered on APAF1, with proteins like SFN, SFN, SERPINB5 (Figure 3D).

**Figure 3.**
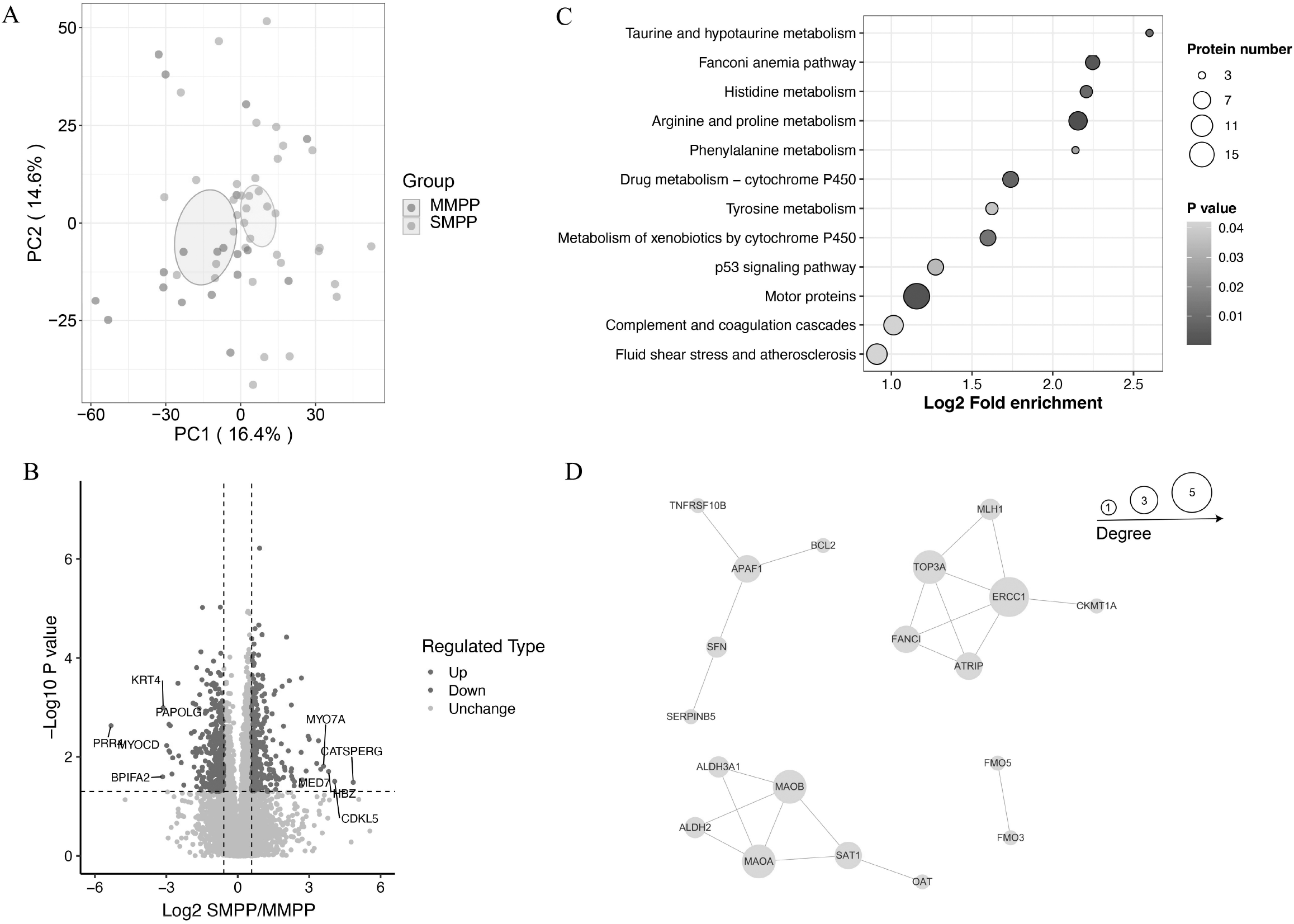
Proteomics Analysis between severe and non-severe patients in BALF. A. The proteomics signatures analysis by Principal Component Analysis (PCA) B. The volcano plot derived from a targeted metabolomic analysis illustrates the top BALF proteomics that were increased (shown in red) or decreased (shown in blue) C. KEGG pathway enrichment analysis of differential proteins. D. Functional network analysis of proteins enriched by KEGG

### BALF metabolite profiles for patients with infection

Metabolomic analysis in 57 patients was conducted to characterize the physiological state of the disease. The OPLS-DA and PCA analysis sets the BALF samples of the patients into the two-dimensional space by a distinction based on severe and mild, indicating differential scores between them with their BALF metabolite profiles (Figure 4A and S3). The differentially expressed metabolites between the BALF of severe and mild groups, identified from the v-fold analysis of such untargeted metabolomic analyses, are respectively viewed as increased (red) and decreased (blue) from the volcano plots (Figure 4B). The KEGG pathway enrichment analysis of metabolites involved in the intersection was performed. The pathways highlighted were enriched with metabolic differences related to inflammation, among others, namely efferocytosis, fatty acid elongation, and D-amino-acid metabolism (Figure 4C). In the coordination of the proteins related to these metabolic pathways, arginine and plasmid metabolism were considered essential (Figure 4D and S4).

**Figure 4.**
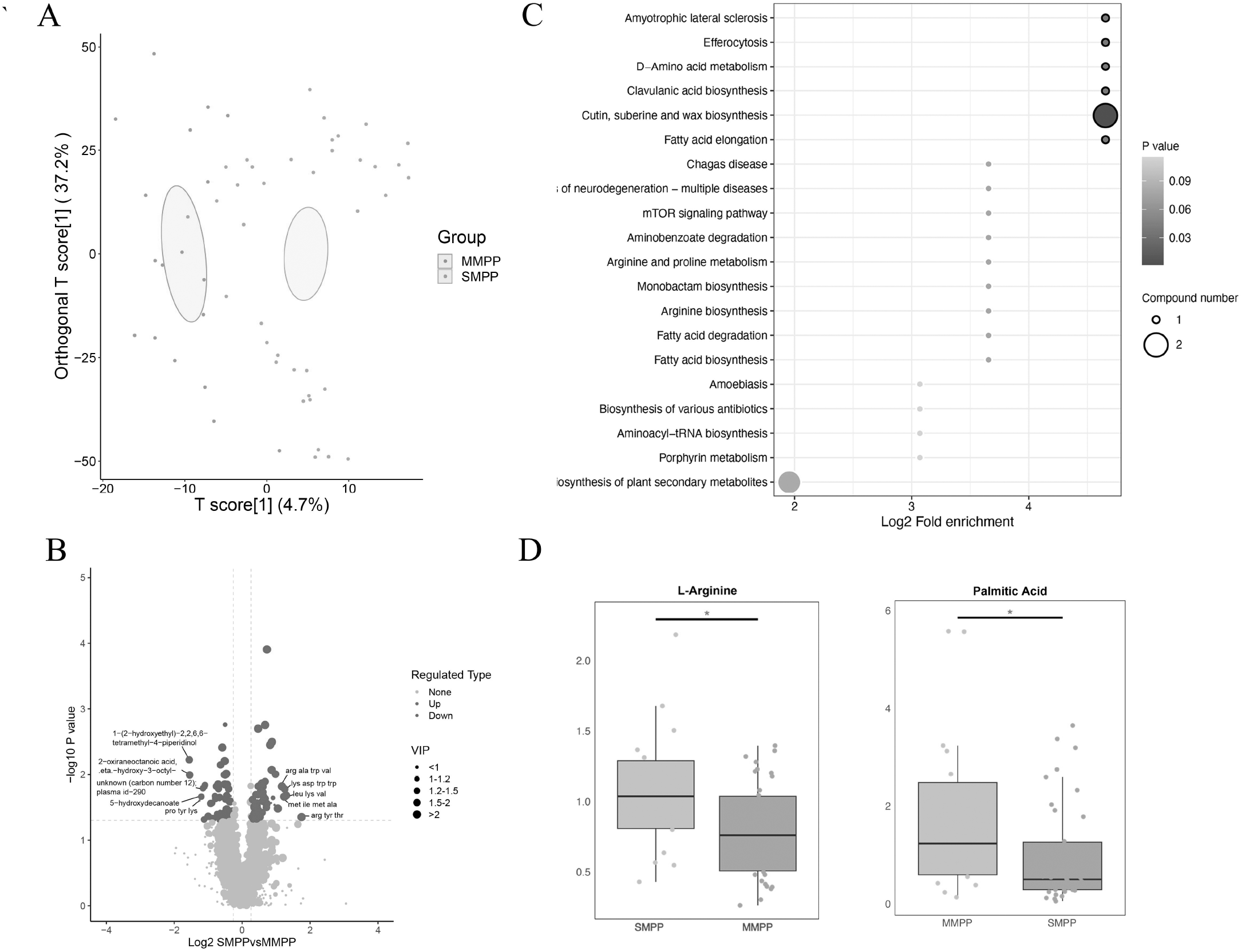
Metabolic Analysis between severe and non-severe patients in BALF. A. The metabolic signatures analysis by Orthogonal Partial Least Squares-Discriminant Analysis (OPLS-DA) B. The volcano plot derived from a targeted metabolomic analysis illustrates the top BALF proteomics that were increased (shown in red) or decreased (shown in blue) C. KEGG pathway enrichment analysis of differential metabolism. D Key metabolism related to KEGG enrichment

### Correlation Analysis and Biomarkers to Predict Patient Mortality

A correlation analysis was conducted on all proteins and metabolites enriched in the KEGG pathway from bronchoalveolar lavage fluid (BALF) samples. In connection with that, it was demonstrated by Figure 5A that there was a large difference and correlation between the two proteins, APAF1 and SERPINB5, enriched in the Fanconi anemia pathway, with the metabolite, L-arginine. When comparing and contrasting APAF1 in the severe to the mild group, it was upregulated, and SERPINB5 and L-arginine were downregulated in the severe to the mild group. In addition, statistically different levels of ERCC1 and MLH1 were correlated with plasmid acid, both downregulated in severe when compared to the mild group and underpinned by the p53 signaling pathway.

**Figure 5.**
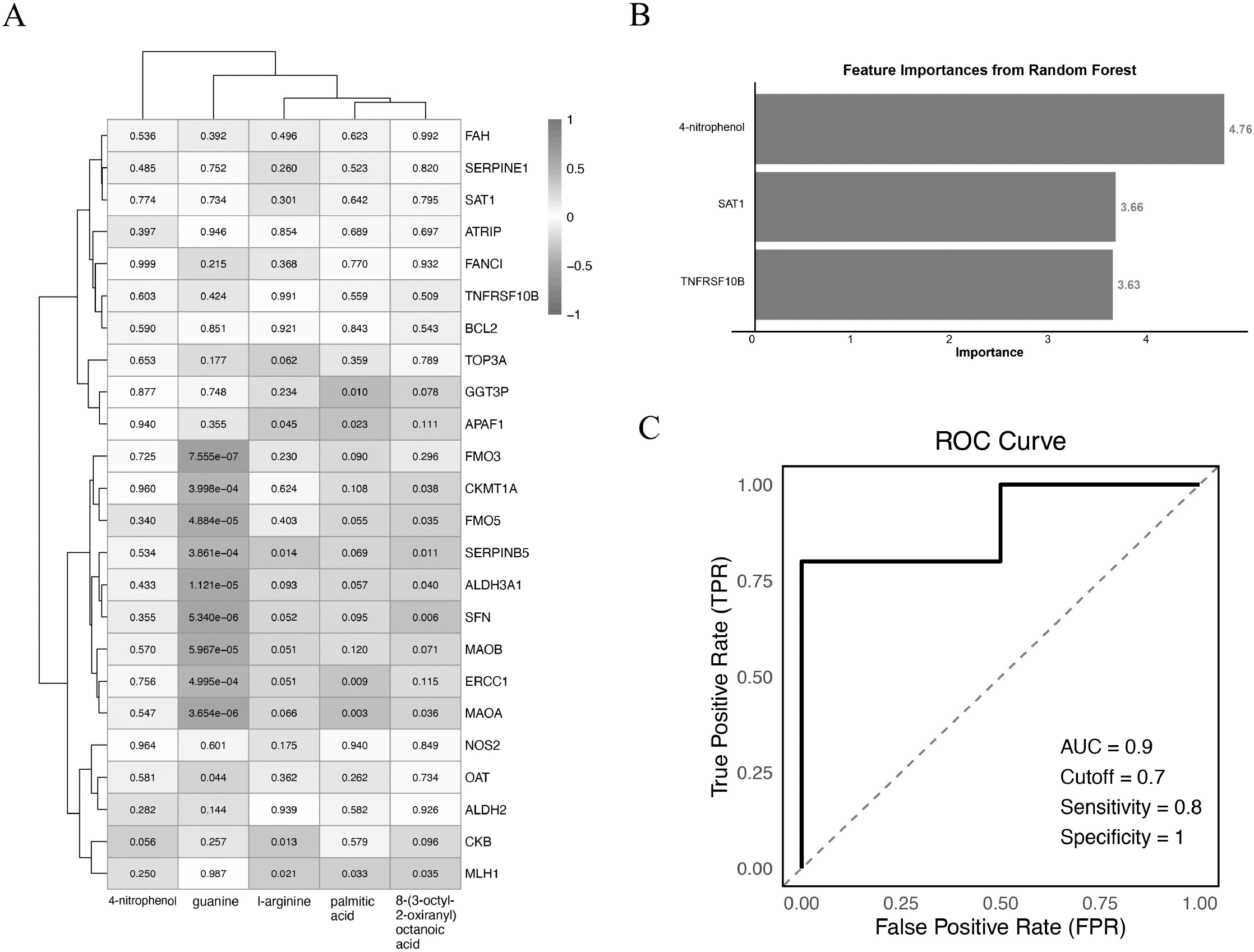
Correlation Analysis and biomarkers. A. KEGG-enriched protein and metabolite association analysis; B. Key biomarker importance identified using a random forest model;C. ROC curve of three biomarkers calculated for the classification of SMPP and MMPP;

Based on the ratio values from SMPP and MMPP groups, the original protein metabolites were arranged in decreasing order. Random forest training was conducted on the highest-ranking protein metabolites, where TNFRSF10B, SAT1, and 4-nitrophenol were revealed to be potential biomarkers (figure 5B). These markers strongly distinguish between SMPP and MMPP. The AUC was simulated to be equal to 0.9, with sensitivity and specificity of 0.8 and 1.0, respectively, as the cutoff value at 0.7 (figure 5C).

## Discussion

Proteins and metabolites in the rapid progression of infectious diseases, including Staphylococcus aureus and COVID-19.^10,17–19^Although early studies have explored the metabolomics of MPP patients’ blood^16^, they have not directly investigated the specific sites of lung infections. We provide the first report on the alveolar lavage fluid sampling in children who had MMP infection; at the same time, we performed proteomics and metabolomics analyses on BALF. We found a high enrichment of differentially expressed proteins and metabolites in inflammatory pathways. Using AI models to train the biomarkers, we highlighted the factors that could mark severe and mild cases effectively.

In the clinical information and examination of 38 severe cases and 19 mild cases, no significant differences were established for age and gender. The levels of PCT and LDH were significantly elevated in the severe group, which concurs with the findings in the article^20–22^. However, no significant differences were observed in white blood cells, lymphocytes, CRP, IL-6, and SAA, which is inconsistent with the article at https://tp.amegroups.org/article/view/131476/html.^22^ The observed discrepancy can presumably be attributed to the fact that white blood cells, lymphocytes, CRP, IL-6, and SAA are not specific indicators of MPP infection and may be influenced by other factors, such as chronic inflammatory diseases, administration of medications and treatments, age, and so on.^23,24^ Hence, the identification of specific proteins and metabolic markers is essential in tackling severe MPP infections.

The article performed PCA analysis of plasma proteomics and found strong separation between plasma proteins in the HC, MMPP, and SMPP groups. It also identified a large number of differentially expressed proteins between the two groups, suggesting that significant differences exist between hospitals and between groups at different disease severity levels. The expression pattern clustering analysis showed that most of the proteins were arranged in a linear manner from healthy through mild to severe stages. Functional enrichment analysis of the intersection of differential proteins showed that they were mainly enriched in the inflammatory pathway and the Complement and coagulation cascades, consistent with previous studies^25,26^. All these lead us to believe that inflammation response and complement activation in plasma really do account for the differences in severity observed in MMP infections.

Thereafter, in order to undertake a better analysis of the pathophysiology of respiratory tract infections and reasons for the severity of Mycoplasma pneumonia disease, we collected alveolar lavage fluid for proteomic and metabolomic analysis. Analyses utilizing principal component analysis and differential analysis indicated that proteins and metabolites in the bronchoalveolar lavage fluid from the Mild and Severe groups had sufficient separation and differences. Functional analysis and PPI network analysis showed that the main differential proteins and metabolites in alveoli were enriched in relevant pathways, including Arginine_and_proline_metabolism, Fanconi anemia, p53 signaling, etc. analysis between proteins and metabolites revealed that proteins involved in inflammatory pathways showed a significant correlation with differential metabolites, indicating a regulatory relationship between proteins and metabolites. For instance, APAF1 and SERPINB5 proteins in p53 signaling were significantly correlated with L-arginine metabolites, while L-arginine metabolites also showed significant association with the Efferocytosis pathway. This indicates that the APAF1 and SERPINB5 proteins may mediate changes in L-arginine metabolites, which would drive changes in the pathway. The protein APAF1 promotes inflammation, which is consistent with the reports in the article.^27,28^ And the proteins L-arginine and SERPINB5 have been reported in the article to alleviate inflammation.^29,30^ In the same way, the proteins ERCC1 and MLH1 in the Fanconi anemia pathway found to be significantly correlated with plasmid acid metabolites, while L-arginine metabolites were related to the Fatty acid elongation pathways. It indicates that the ERCC1 and MLH1 proteins in the Fanconi anemia pathway may mediate changes in plasmid acid metabolites that drive changes in the Fatty acid elongation pathway. There are research reports indicating that ERCC1, MLH1, and plasmid acid all have anti-inflammatory effects.^31–34^

At present, there are no biomarkers available for clinically assessing Mycoplasma pneumonia (MPP) infection severity: This hinders clinicians’ treatment and disease progression decision-making. The lack of reliable markers, in turn, makes it highly difficult to identify these high-risk candidates who need intensive interventions. This limitation amplifies the need for accurate and easily detectable biomarkers that can be of assistance in giving timely information about infection and its severity. In the study, we tackled this question by relying on the key protein metabolites that were enriched through functional analyses. After determining a set of key metabolites, we ranked them according to their association disease severity. Finally, we trained the classifier using a random forest model based solely on three selected biomarkers (TNFRSF10B, SAT1, and 4-nitrophenol). A strong and performant model was built using this approach. When applied to the test dataset, the classifier achieved strong performance, with Area Under the Curve of 0.9, specificity of 1.0, and sensitivity of 0.8. These findings indicate that this diagnostic tool has significant potential for accurate and reliable application.The results suggest that these three markers could conceivably act as important grading criteria for an MPP infection, enabling physicians to assess patients better and direct treatment approaches accordingly.

### Limitations of the study

The limitation of this research is a single-center study, and the data were collected exclusively Huazhong University of Science and Technology Union Shenzhen Hospital. A larger population of children needs to be included. Also, the research did not collect bronchoalveolar lavage fluid from healthy children due to this procedure being invasive, hence it lacked an analysis of pulmonary metabolism and proteomics in healthy children.

## Conclusion

This research has great potential to contribute to both clinical care and understanding of MPP infections in children. By using integrative multi-omics approaches (proteomics, metabolomics) and making informed use of machine learning, it identified specific biomarkers associated with infection severity, predicting better, along with personalized treatment and improved disease monitoring opportunities. An in-depth understanding of the host immune response precipitated by this study provides the roadmap to novel targets and preventive strategies not seen before. Ultimately, this drives us toward a more principled precision medicine approach to enhance diagnostics and therapy for pediatric patients

## Methods

### Patients

The patients and controls were recruited from the Huazhong University of Science and Technology Union Shenzhen Hospital from August 2023 to August 2024. Diagnosis of pediatric mycoplasma pneumonia was made based on guidance from the Chinese Medical Association as follows: (i) fever and acute respiratory symptoms (e.g., cough, tachypnea, or difficulty breathing), or any combination of the two, in children between the ages of 3 and 15, regardless of sex; (ii) low breathing or dry, wet rales; (iii) chest X-ray revealing enlarging hilar lymph nodes, lung gate shadows, bronchopneumonia and interstitial pneumonia, and large, high-density shadows; (iv) PCR or metagenomic next-generation sequencing-positive. Exclusion criteria: (i) pneumonia history within 3 months; (ii) history of preterm birth, asthma, or severe cardiovascular conditions such as congenital heart disease, cardiomyopathy, liver or kidney; (iii) family history of tonsillar hypertrophy; (iv) the investigator’s opinion towards unsuitability for participation in the study.

Severe MPP was defined by the presence of one or more of the following criteria: (1) persistent high fever (≥39°C) lasting for ≥5 days; or fever lasting for ≥7 days with no decline in peak temperature; (2) clinical findings suggestive of severe illness that continue associations with bronchial plasticity, asthma exacerbation, pleural effusion, or pulmonary embolism, such as wheezing, shortness of breath, respiratory distress, chest pain, or hemoptysis; (3) the presence of extra-pulmonary complications but not meeting critical condition criteria; (4) oxygen saturation (SpO2) ≤93% with room air at rest; (5) imaging findings that include: (a) more than 2/3 of a single lung lobe affected with uniform high-density consolidation, or high-density consolidation in two or more lung lobes (regardless of the extent of the affected area) may be associated with moderate to large pleural effusion or localized bronchitis; (b) diffuse bronchitis in more than 4/5 of a single lung or bilateral lung lobes, possibly with bronchial inflammation, mucus plug formation, and atelectasis; (6) a steadily worsening clinical picture, with imaging showing a disease progression of more than 50% within 24-48 hours. The Ethics Committee of Huazhong University of Science and Technology Union Shenzhen Hospital approved this study. The protocols for this research study were carried out according to the guidelines set forth by the Ethics Committee.

### Sample Collection

Bronchoalveolar lavage fluid (BALF) volumes collected were ≥8 ml and the sample was centrifuged at 10 minutes for 12,000 g to collect the pellet. The pellet was resuspended 1-2 times in sterile 1x PBS and was centrifuged again at 12,000 g for 10 minutes to collect the pellet. Additionally, 3 ml of venous blood was drawn into an EDTA-K2 anticoagulant tube, thoroughly mixed by inverting the tube 5-8 times and centrifuged for 10 minutes at 1,300 rpm to separate the plasma. At least 300 μL of plasma was transferred to a 1.5 ml EP tube and immediately stored at −80°C for further analysis.

### Protein Extraction and Analysis

Sample collected from alveolar tissues was taken from −80°C and mixed with a 4x volume of lysis buffer. After heating, sonicate the samples. The amounts of protein digested were equal. After an aliquot of the lysis buffer was added to control for volumes, one volume of prechilled acetone was added. Trypsin was added at a 1:50 ratio (w/w, protease) for overnight digestion. 5 mM DTT was added. Iodoacetamide to a final concentration of 11 mM was then added, after which the samples were incubated for 15 minutes at room temperature, and protected from light. The peptides were further dissolved in mobile phase A and separated using the EASY-nLC 1200 ultra-high-performance liquid chromatography system. Mobile phase B consisted of 0.1% formic acid and 90% acetonitrile. After separation, the peptides were injected into the nano-spray ionization ion source for ionization and analyzed on the Orbitrap Exploris 480 mass spectrometer. Both precursor and fragment ions were detected and analyzed using a high-resolution Orbitrap system.

### Metabolite Sample Preparation and Analysis

250 µL of the original sample was treated with 4 volumes of the extraction buffer. Then, with vigorous vortexing, the sample was sonicated. Thereafter, a further 50 µL of ACN was added, and the sample was fused again by sonication. Metabolites were separated using a Waters UPLC ultra-high-pressure liquid chromatographic system coupled with a Waters ACQUITY UPLC BEH C18 Column (1.7 µm, 2.1 mm × 100 mm). The injection volume was 10 µL and eluted at a flow rate of 400 µL/min at a column temperature of 45°C. Mobile phase A was an aqueous solution containing 0.1% formic acid, while mobile phase B was acetonitrile containing 0.1% formic acid. The UPLC-separated metabolites were then introduced into the ESI ion source for ionization and subsequently analyzed by the Tims TOF Pro mass spectrometer.

### Bioinformatics Analysis

The quantitative metabolite data were further processed by using data filtration after a thorough database matching had been done. Then fold change was determined by taking into account the corrected expression levels between the two groups using samples that were subject to repetition. In addition, the univariate T-test P-values and multivariate analysis. PCA and OPLS-DA included accounted for the computation of the VIP scores.

This approach assists in determining which proteomes and metabolites are significantly different. Further, bioinformatics analyses and functional analyses on these differential metabolites were performed. When the samples were studied with three or more sample groups, P-values by ANOVA analyses across the groups were obtained and differential metabolites were screened from those P-values. The proteomes and metabolites were clustered with respect to their expression, followed by Gene Ontology analysis (GO) and Kyoto Encyclopedia of Genes and Genomes (KEGG) enrichment analysis.

### Biomarker selection of MPP

The data is integrated with the alveolar differential proteome and metabolites, and, according to the sample grouping information (MMPP and SMPP), the data is divided into a training set (data train) and a test set (data test) with a ratio of 7:3. A random forest model is built upon the training set feature data to predict the sample group labels. The random forest function builds the model while discrepancy analysis on features is done to feature importance using the importance function for the trained random forest model. The features are ranked according to their importance values in a descending manner, and their importance bar chart is constructed. Then, the model is validated on the test set (data test) and the probability of each sample belonging to a class is obtained. Based on these values of predicted probabilities, the ROC curve is constructed, and then the AUC is formed.

### Resource avaiability

Lead contact Further information and requests for resources and reagents should bedirected to and will be fulfilled by the lead contact, Chunlan Xiu.

### Materials availability

All unique models generated in this study are available from the lead contactwith a completed materials transfer agreement.

## Acknowledgments

We are particularly grateful to all the people who have given us help with our article.

## Author contribution

Guiqiu Li and Xiulan Lai conceptualized and designed the experiments. Zhili Hu collected and analyzed data. Guiqiu Li and Wenzheng Wang drafted the initial manuscript. Xiulan Lai contributed to data analysis and interpretation, and revisions to the manuscript. Qiaowen Yang provided theoretical support, Lili He and Yun Gaoconducted literature reviews, performed discussion analysis, and also revised manuscript revision. All authors reviewed and approved the final submitted version.

## Ethical Approval

The study was conducted following the Declaration of Helsinki (revised in 2013). The study was approved by the Ethics Committee of Huazhong University of Science and Technology Union Shenzhen Hospital. (Shenzhen, China).

## Data availability

All data supporting the finding of this study are provided within the paper and the Supplementary Information.

## Declaration of Generative AI and AI-assisted technologies in the writing process

During the preparation of this work the authors used Chatgpt (Website: https://chatgpt.com/) in order to write clearly, coherently, and concisely in the manuscript. After using this tool, the authors reviewed and edited the content as needed and take full responsibility for the content of the publication.

---

**Table 1.**
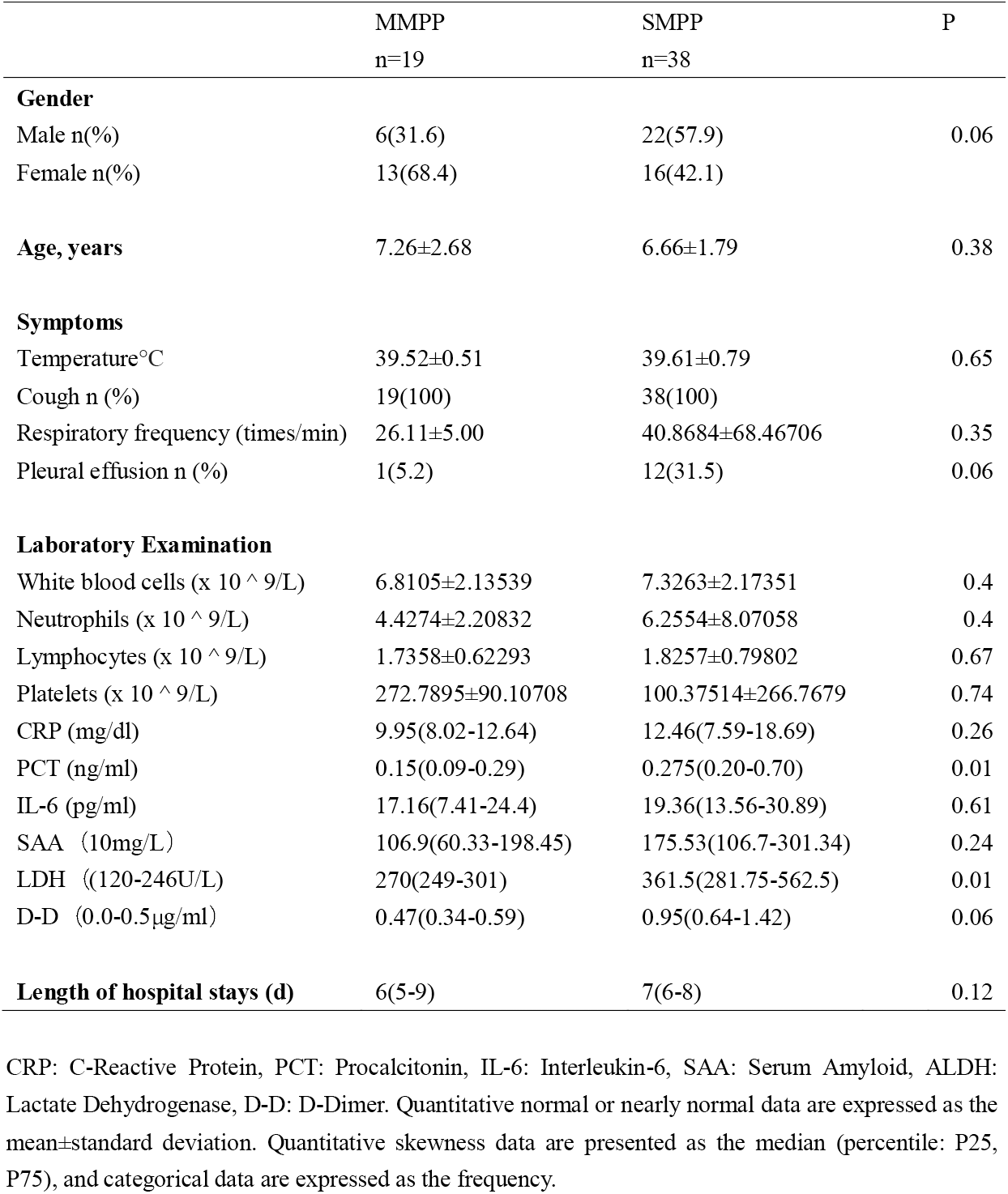
Demographic characteristics of children between MMPP group and the SMPP group.

