## Supplementary material for "Multi-Omics and Machine Learning-Based Profiling of Severity Signatures in *Mycoplasma Pneumoniae* Infection in Children": supplmental figures

*Running title: Exploring Mycoplasma Pneumoniae Infection by Multi-Omics*


Guiqiu Li[2], Wenzheng Wang,[2] Zhili Hu[1], Qiaowen Yang[1], Lili He[1], Yun Gao[1], Xiulan Lai[1]

1. Pediatrics Department, Shenzhen Nanshan District People's Hospital, Shenzhen, 518052, China
2. Clinical Laboratory Department, Shenzhen Nanshan District People's Hospital, Shenzhen, 518052, China

Corresponding author：Xiulan Lai

Phone number：15112662044


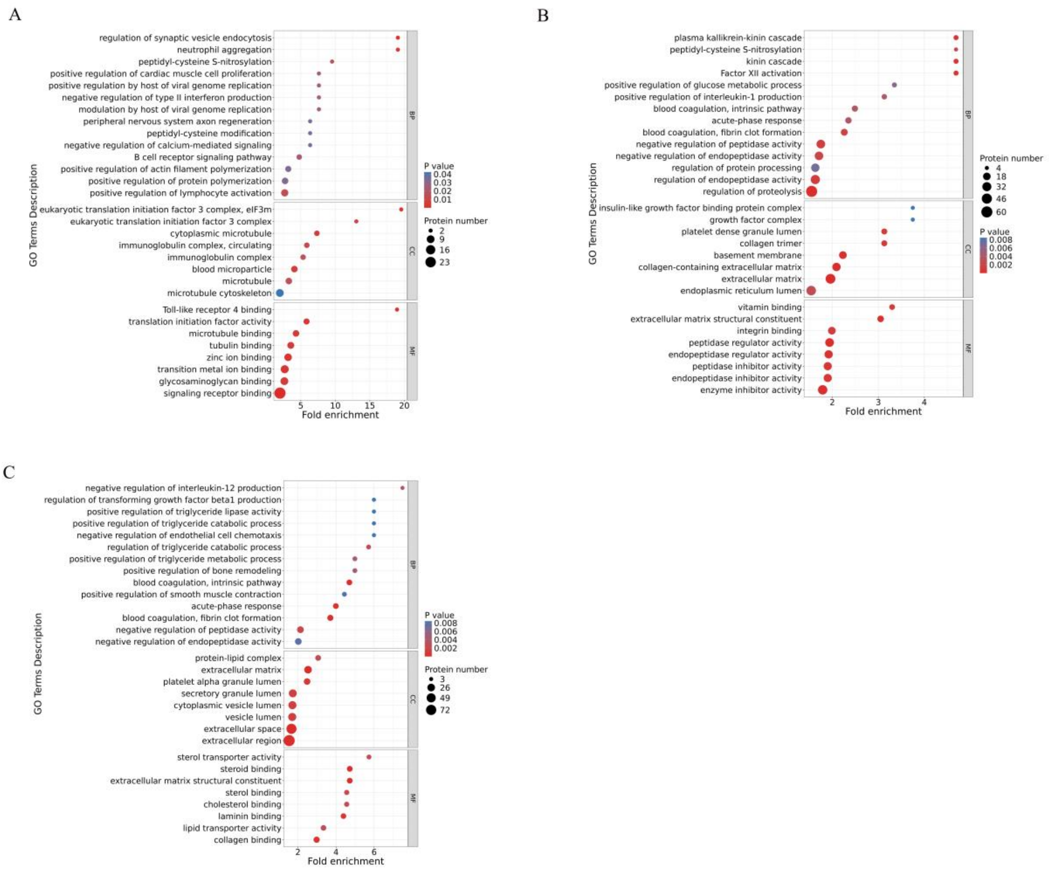


S1. Gene ontology enrichment analysis of differentially expressed proteins in blood sample A SMPP vs MMPP, B SMPP vs HC, C MMPP vs HC

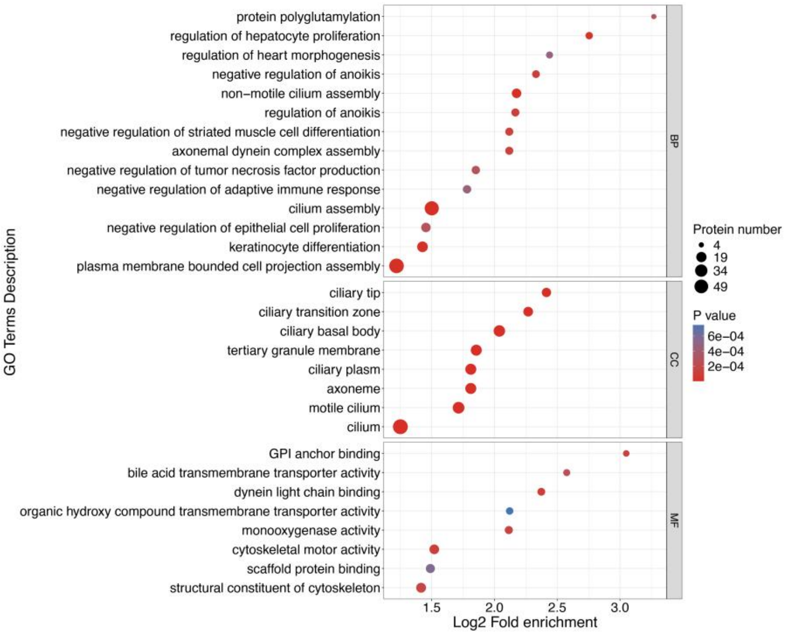


S2. Gene ontology enrichment analysis of differentially expressed proteins in BAIF sample

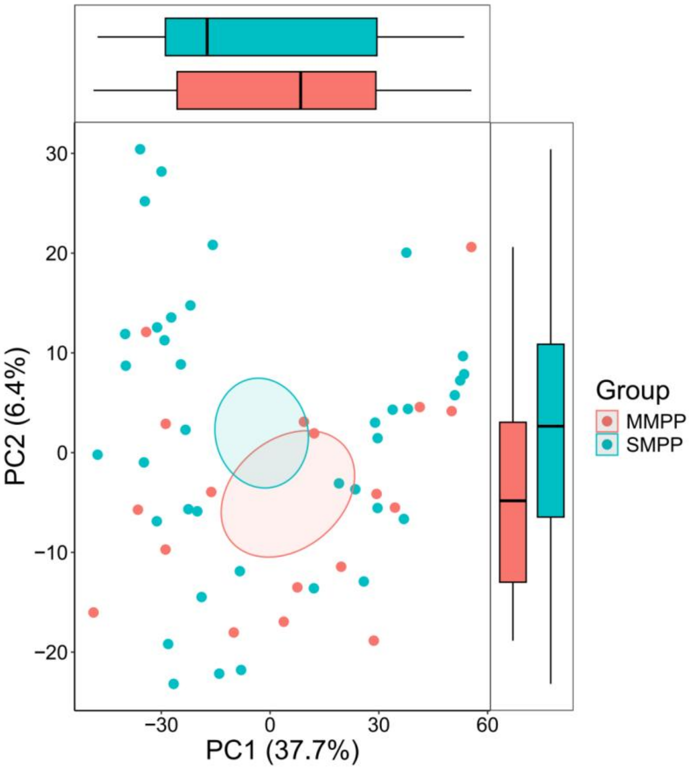


S3. Alveolar Metabolite Principal Component Analysis (PCA Plot). Red represents MMPP, and green represents SMPP.

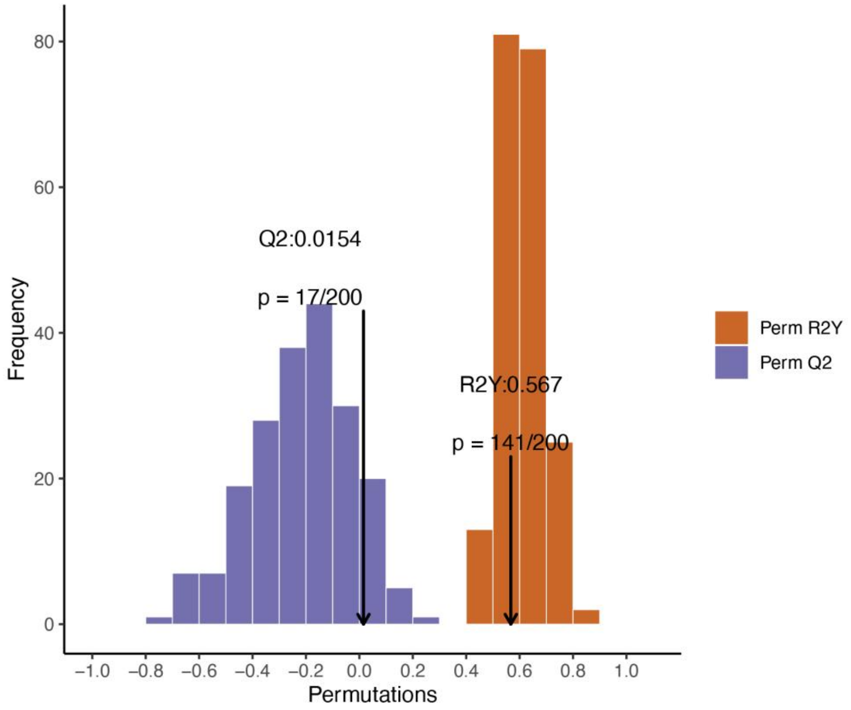


S4. The OPLS-DA model validation: The x-axis represents the accuracy of the model, and the y-axis represents the frequency of the model's accuracy.

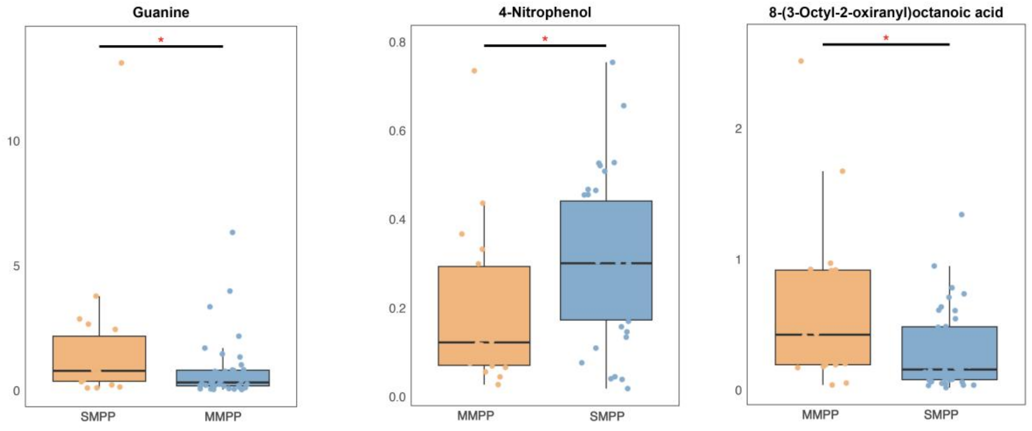


S5. Key metabolites enriched in the alveolar metabolome according to KEGG pathway analysis.
